# Genetic Analysis of Osteoblast Activity Identifies *Zbtb40* as a Regulator of Osteoblast Activity and Bone Mass

**DOI:** 10.1101/828525

**Authors:** Madison L. Doolittle, Gina M Calabrese, Larry D. Mesner, Dana A. Godfrey, Robert D. Maynard, Cheryl L. Ackert-Bicknell, Charles R. Farber

## Abstract

Osteoporosis is a genetic disease characterized by progressive reductions in bone mineral density (BMD) leading to an increased risk of fracture. Over the last decade, genome-wide association studies (GWASs) have identified over 1000 associations for BMD. However, as a phenotype BMD is challenging as bone is a multicellular tissue affected by both local and systemic physiology. Here, we focused on a single component of BMD, osteoblast-mediated bone formation in mice, and identified associations influencing osteoblast activity on mouse Chromosomes (Chrs) 1, 4, and 17. The locus on Chr. 4 was in an intergenic region between *Wnt4* and *Zbtb40*, homologous to a locus for BMD in humans. We tested both *Wnt4* and *Zbtb40* for a role in osteoblast activity and BMD. Knockdown of *Zbtb40*, but not *Wnt4*, in osteoblasts drastically reduced mineralization. Additionally, loss-of-function mouse models for both genes exhibited reduced BMD. Our results highlight that investigating the genetic basis of *in vitro* osteoblast mineralization can be used to identify genes impacting bone formation and BMD.

## INTRODUCTION

Osteoporosis is a metabolic disease characterized by progressive bone loss leading to skeletal fragility and fracture ^1^. Approximately 200 million people worldwide have or are at risk of developing osteoporosis ^2^, leading to ∼8.6 million fractures annually ^3^. As the proportion of aged persons worldwide is increasing, osteoporosis is becoming an even greater public health burden ^4^. Importantly, one in three women and one in five men will suffer an osteoporotic fracture ^5-7^. Family history remains the strongest risk factor for development of osteoporosis and studies in animal models reinforce that this is a complex genetic disease ^8-10^. Furthermore, fracture-related traits, such as bone mineral density (BMD), are among the most heritable disease associated quantitative traits (h^2^>0.50) ^10-12^. Thus, increasing our understanding of the genes influencing osteoporosis is critical for the development of approaches for its treatment and prevention.

Bone is not a static tissue, but rather it is constantly changing to adapt to the current needs of the organism. Turnover in bone is accomplished by three main cell types: the *osteoblasts* which are responsible for forming new bone, the *osteoclasts*, which are the cells responsible for bone resorption, and the *osteocytes* which orchestrate bone turnover through paracrine regulation of both the aforementioned cell types ^13-15^. During normal remodeling, bone formation equals resorption and net bone mass is constant. When remodeling is imbalanced, such as occurs in osteoporosis, bone resorption exceeds formation leading to a net loss of bone and a decrease in BMD. Primary imbalances in bone remodeling occur due to hypogonadism, such as in menopause in women ^16^, and with age ^17^. Secondary bone remodeling imbalances can result from numerous conditions which alter systemic processes such as hormone regulation, or nutritional intake and absorption. These conditions include, but are not limited to, Type II Diabetes ^18^, inflammatory bowel disease ^19^, anorexia nervosa ^20^, chronic kidney disease ^21^. Furthermore, several environmental factors such as smoking ^22^, air pollution ^23^, alcohol consumption ^24^, high fat diets ^25^ and use of common medications such as proton pump inhibitors ^26^, barbiturates ^27^, glucocorticoids ^28^, and NSAIDS ^29^ are associated with reduced bone mass and/or fracture incidence. Also, bone mass is influenced by genetic signals that emanate from a plethora of organ systems ^30^. While it is important to identify all factors that influence bone mass, the complexity of BMD means that any individual genetic association may exert its influence through many potential mechanisms, confounding our understanding of the associated biology.

Genome-wide association studies (GWAS) have been extremely successful in identifying loci affecting osteoporosis-related traits. These studies have primarily focused on BMD and surrogates for BMD, such as estimated BMD (eBMD) measured using ultrasound of the calcaneus ^31^. To date, the largest GWAS for eBMD identified 1103 independent associations, collectively explaining ∼20% of the variance for this phenotype ^32^, emphasizing the idea that much remains to be identified. Additionally, mouse mapping studies have identified large numbers of quantitative trait loci (QTL) ^8^ and genome-wide associations for bone traits ^33^. While it is imperative that all genetic influences on osteoporosis be identified, there is an urgent need for genetic studies that fill in the holes in our knowledge that arise from complex nature of the BMD phenotype.

In this study, we focused on a specific component of bone remodeling, and performed the first cell-specific genetic analysis of osteoblast function. The advantage of this strategy is that genetic regulation of osteoblast activity is, in theory, simpler than the genetic regulation of bone mass as the environment can be more controlled and the read-out is based on the intrinsic action of a single cell type. Therefore, identified associations will provide a clearer path to go from locus to gene to mechanism, and it provides a way to focus on the component of bone remodeling that is in the most need for new anabolic therapeutic targets.

## RESULTS

### Genetic Analysis of Osteoblast Activity

#### Measuring Osteoblast Activity in a Panel of Inbred Mouse Strains

One of the primary goals of this study was to “simplify” the genetic analysis of complex skeletal traits by focusing on a cellular phenotype. Primary osteoblasts isolated from neonatal calvariae, when cultured in osteogenic media, form “nodules” of mineralization that are similar to the mineralized matrix found in bone and *in vitro* mineralization directly reflects the primary activity of bone forming osteoblasts (42). We began by isolating primary calvarial osteoblasts from 22 inbred strains of mice, differentiated them into mineralizing osteoblasts, and measured the amount of mineral after 10 days in culture. We observed high levels of inter-strain variation in mineralization (**Fig 1a**), with a 3.9-fold increase between the strain with the lowest (BALB/cJ) and highest (C3H/HeJ) mineralization values. The broad sense heritability (H^2^) of mineralization was 0.87 (P<2.2 × 10^−16^).

**Figure 1:**
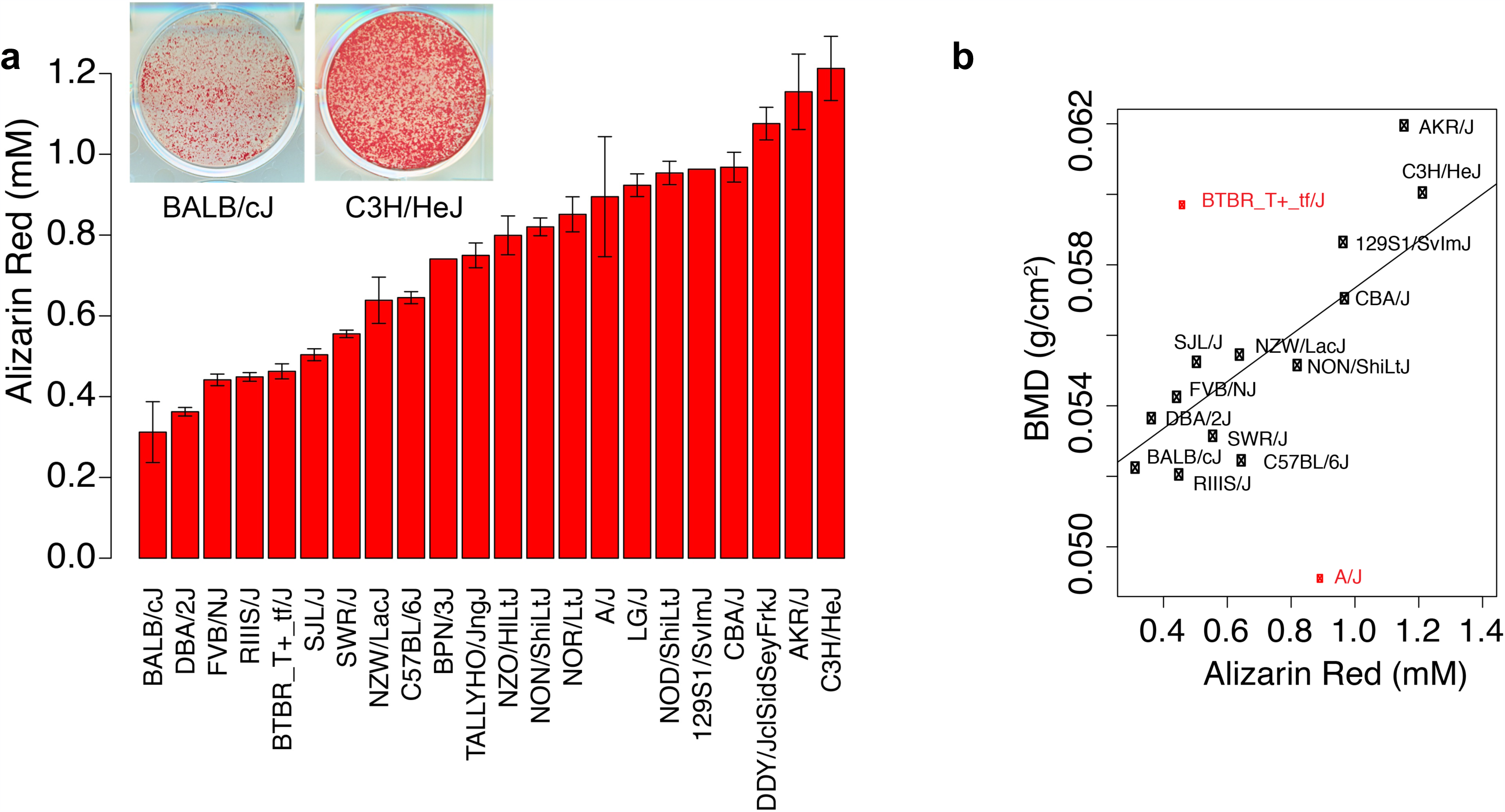
*In vitro* osteoblast mineralized nodule formation is a heritable trait and correlates with *in vivo* BMD. (A) Quantification of Alizarin Red-stained cultures of calvarial osteoblasts isolated from 22 inbred mouse strains after 10 days of osteogenic differentiation. Data are presented as mean ± S.E.M. (B) Alizarin Red values correlate significantly with areal BMD in mouse strains (*r*^*2*^=0.54 *p=0.03, r*^*2*^= 0.89 *p=3.3*×*10*^*-5*^ excluding outliers, BMD data from Ackert1^34^, https://phenome.jax.org/)

We previously generated total body BMD data for 16 of the 22 strains (http://phenome.jax.org, Ackert1 dataset) ^34^. As would be expected, mineralization and BMD were positively correlated (r=0.54; P=0.03) (Fig 1b). These data indicate that *in vitro* osteoblast mineralization is highly heritable and confirm that variation in osteoblast activity is an important contributor to BMD.

#### Genome-Wide Association Study for Osteoblast Activity

We next sought to identify genome-wide associations influencing osteoblast activity. To accomplish this, associations between mineralization and ∼196K SNPs across the 22 strains were calculated using the Efficient Mixed Model Algorithm (EMMA, ^35^). Three loci on Chromosomes (Chrs) 1, 4, and 17 exceeded the permutation-derived significance threshold of P=2.5 × 10^−6^ (**Fig 2a**). The three associations were independent (r^2^ among all top SNPs <0.25). Osteoblasts from strains homozygous for the reference (C57BL/6J) allele on the Chrs. 1 and 17 associations were less active, whereas osteoblasts from strains homozygous for the reference allele on Chr. 4 were more active (**Fig 2b,c,d**). Mouse QTL for BMD and other bone traits have been found overlapping these associations, providing further support that they are true associations ^8^.

**Figure 2:**
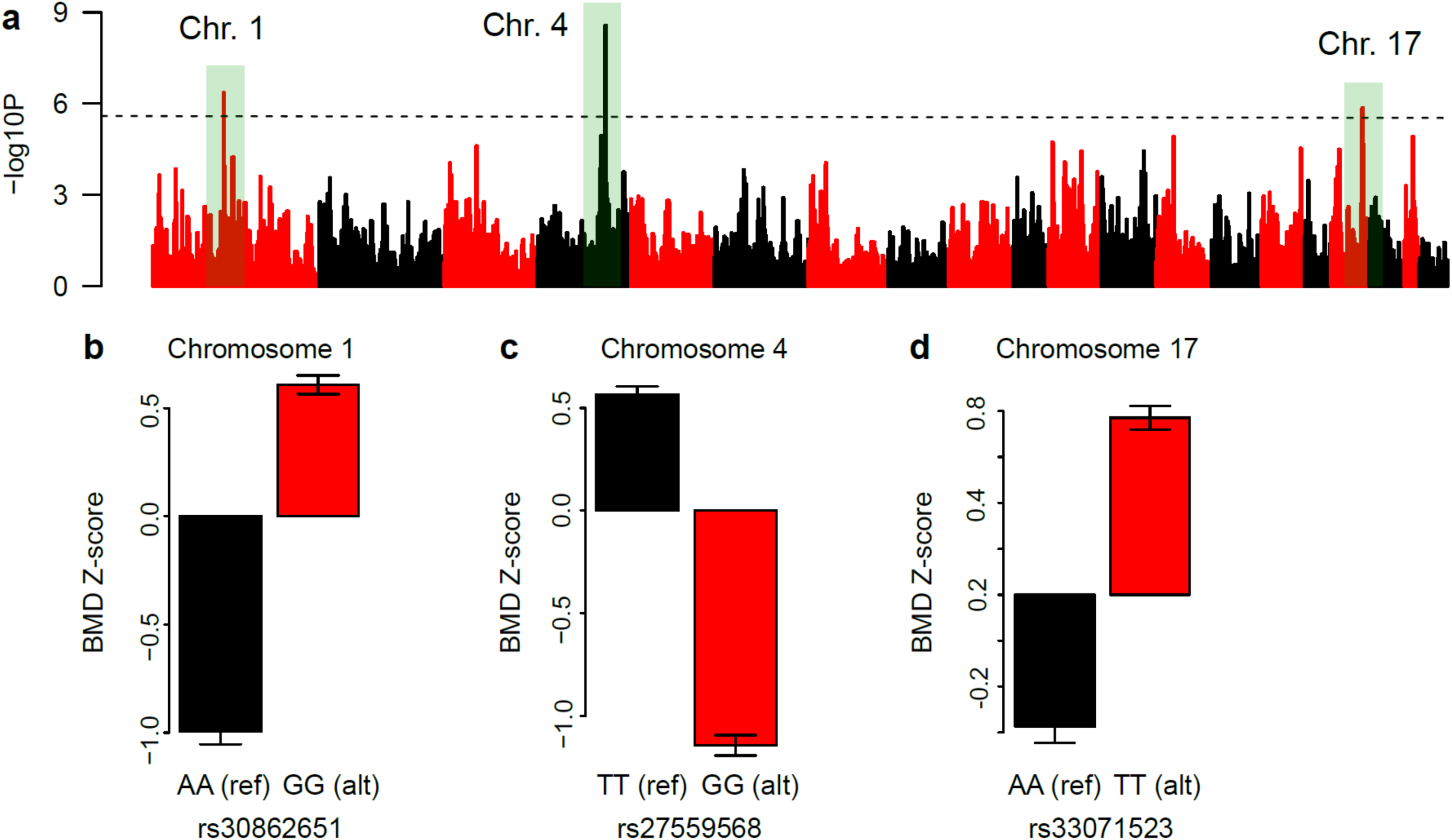
Genome-wide association analysis identifies three loci for osteoblast activity. (A) Association results (-log10 of the P-value) for 196,330 SNPs and Alizarin Red quantification values across 22 inbred mouse strains. The dotted black line represents the genome-wide significant threshold. (B) Effect plots for the lead SNPs at each of the three significant loci. Ref=refence (B6) allele; alt=non-reference allele.

#### Characterization of GWAS loci

For the purpose of characterizing each locus, we conservatively defined the location of each association as the region spanning the most significantly associated haplotype flanked by 500 Kbp upstream and downstream. Based on this definition, the associations spanned 1.0 (Chr. 1), 1.2 (Chr. 4), and 2.1 Mbp (Chr. 17). We first evaluated whether any of the most significantly associated SNPs or SNPs in high LD (r2>0.8) were potentially functional (missense, frameshift, etc.). There were no genes, however, across the three regions that harbored such a variant. We then used two expression datasets to prioritize genes: RNA-seq time-course profiles (days 0-18, every 2 days) from FACs sorted calvarial osteoblasts isolated from C57BL/6J mice (GSE54461, ^36^) and local eQTL in demarrowed cortical bone (GSE27483, PMIDs 21490954 and 23300464) in the Hybrid Mouse Diversity Panel (HMDP, a set of 100 inbred strains, including 13 of the strains used in our pilot ^37^). Of the genes expressed in osteoblasts across differentiation (as determined using the RNA-seq data), *Jarid1b, 9530009M10Rik*, and *Ptprv* on Chr 1 and *Zbtb40, A430061O12Rik, 2810405F17Rik, Rap1gap*, and *Ece1* on Chr 4 were regulated by local eQTL (P<1.0 × 10^−6^) in cortical bone (Supplemental Table 1). However, of these, only the lead SNP for the *Ptprv* eQTL on Chr 1 was in strong linkage disequilibrium (r^2^>0.8) with the lead SNP for the mineralization association. Specifically, the lead SNP, rs3086265, was the same for both associations and the allele of rs30862651 associated with lower mineralization, was associated with lower levels of *Ptprv* in bone and this is consistent with the known role of *Ptprv* in osteoblast activity ^38,39^. Together these data suggested that the *Ptprv* local eQTL was responsible for the Chr 1 association.

#### Mineralization associations overlap with human BMD loci

We next queried the human regions syntenic with the mouse associations for evidence of association with BMD. To accomplish this, we used data from the largest GWAS for heel estimated BMD (eBMD) performed to date (N∼426K individuals) using data from the UK BioBank ^32^. All three human regions syntenic with the mouse associations harbored at least one genome-wide significant association (**Fig. 3**). The human region syntenic with the mouse Chr. 1 mineralization locus harbored an association influencing eBMD overlapping *LGR6* and *PTPRV* (**Fig. 3a**). The human region syntenic with the mouse Chr. 4 mineralization locus contained multiple independent associations, most of which were centered in the intergenic region between *ZBTB40* and *WNT4* in both species (**Fig 3b**). We also identified two human eBMD associations in the region syntenic with the mouse Chr. 17 mineralization locus, one centered over *PKDCC* and the other overlapping the *ABCG5/8* cluster (**Fig 3c**).

**Figure 3:**
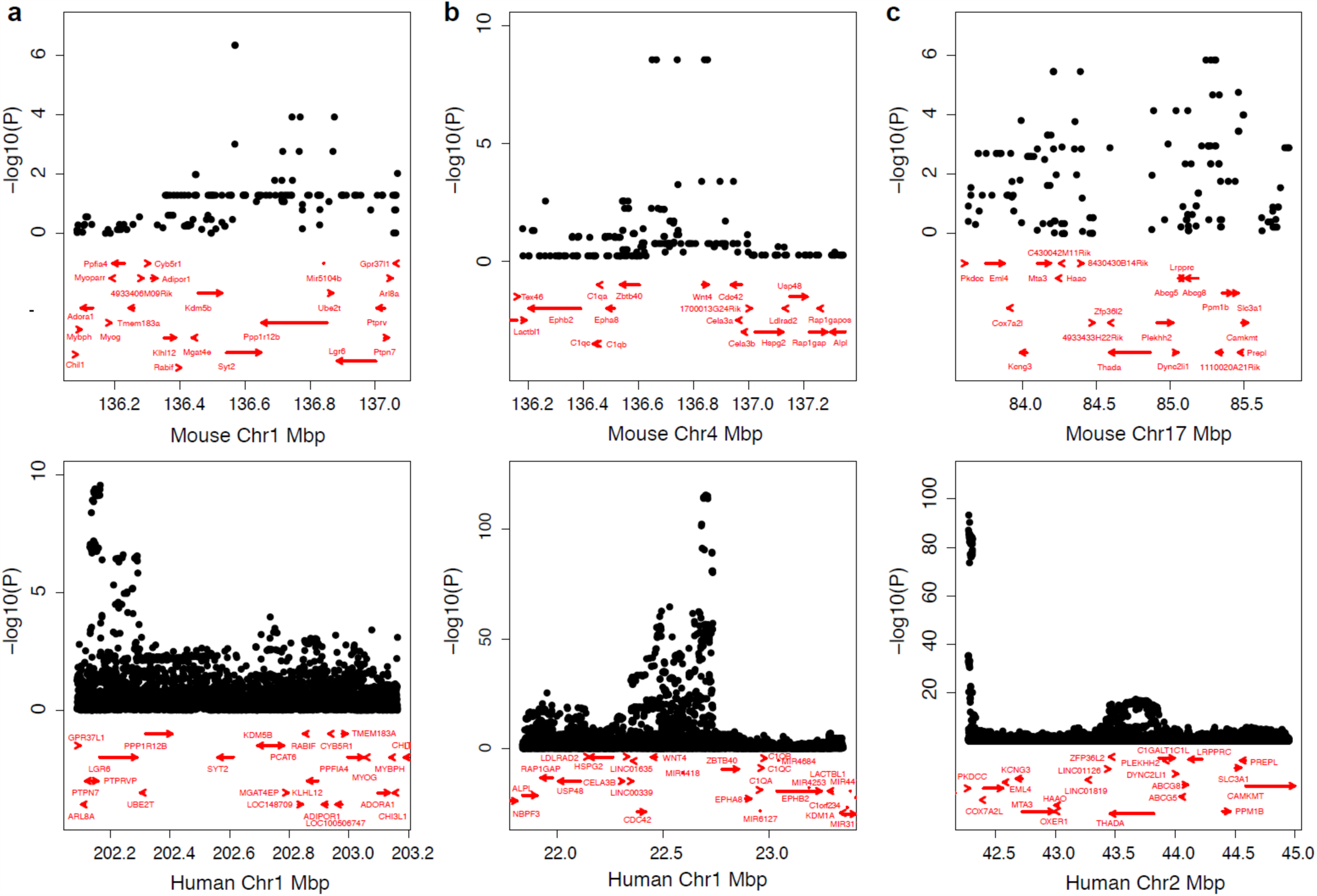
Human regions syntenic with mouse associations harbor genome-wide significant associations for BMD. Plots of GWAS results for eBMD (ref) for the human genomic regions syntenic with the three osteoblast mineralization loci on Chrs. 1 (A), 4 (B), and 17 (C). Note the orientation is reversed in humans for the mouse associations on Chrs. 1 and 4.

### Validation of Candidate Genes Flanking Chr. 4 Locus

As the *ZBTB40 -WNT4* human region is one of the most commonly associated loci with both BMD and fracture in GWAS ^31,32,40-44^, the association in the mouse syntenic region provided grounds for further investigation. Therefore, based on their proximity to the association, we predicted that either *Wnt4* or *Zbtb40* was the causal gene underlying the association for this Chr. 4 locus and sought to experimentally validate the putative roles of these two genes in osteoblast mediated mineralization of the bone matrix.

#### Knockdown of Zbtb40, but not Wnt4, disrupts in vitro osteoblast mineralized matrix formation

To investigate the effect of *Wnt4* knockdown on mineralization, we isolated primary calvarial osteoblasts from mice with *Wnt4 fl/fl, wt/fl*, and *wt/wt* genotypes. To delete *Wnt4*, the cells were then transfected *in vitro* with cre-expressing or empty control plasmids. Using this approach, approximately 38% (P<0.001) and 20% (P<0.001) of the *Wnt4* floxed alleles were excised in *fl/fl* and *wt/fl* cells, respectively (**Fig 4a**). We then quantified mineralized nodule formation at 10 days post-differentiation by Alizarin Red staining and found no significant difference in mineralization as a function of *Wnt4* genotype (*P=0.2077; one-way ANOVA*) (**Fig 4b-c**).

**Figure 4:**
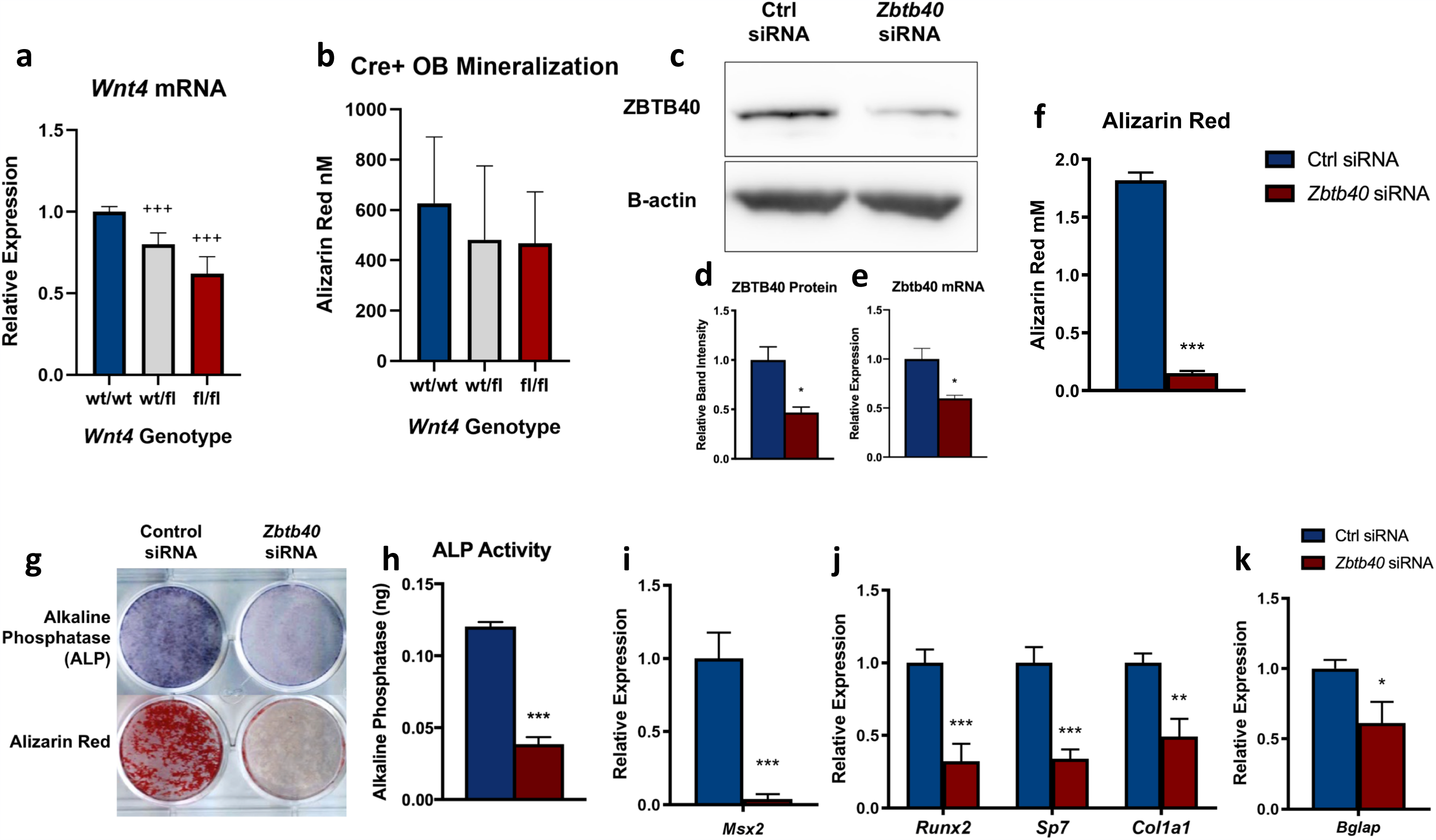
Knockdown of Zbtb40, not Wnt4, inhibits osteoblast differentiation and mineralization: (A) Relative expression of *Wnt4* mRNA levels in calvarial osteoblasts from differing genotypes isolated and transfected with a cre-expressing plasmid *in vitro.* (B) Cell monolayer images and (C) quantification of Alizarin Red-stained cultures of *Wnt4* knockdown (wt/wt n=12, wt/fl n=45, fl/fl n=21). (D) Immunoblot image and (E) quantification of ZBTB40 protein levels in MC3T3 preosteoblast cells indicating knockdown. (F) Relative expression of *Zbtb40* mRNA levels measured by qRT-PCR. (G) Quantification of Alizarin Red-stained Control (Ctrl) or *Zbtb40* siRNA cultures. (H) Cell monolayer images of siRNA-treated cultures stained for Alkaline Phosphatase (ALP) or Alizarin Red measured at day 4 and day 21, respectively. (I) Quantified ALP per well measured by colorimetric activity assay. (J) Relative Expression of mRNA levels of Msx2 at day 0. Relative mRNA levels of early (K) and late (L) osteogenic factors measured at day 4 and day 21, respectively. n=3 independently performed experiments. +++ *= p<0.0001 One-way ANOVA -* =p<0.05 * * =p<0.01 * * * =p<0.001 -*Unpaired T test

As *Wnt4* loss-of-function appeared to have no intrinsic effect on osteoblast mineralization, we next looked to *Zbtb40*. For functional studies, siRNA knockdown of *Zbtb40* was performed in MC3T3-E1 Subclone 4 pre-osteoblast cells ^45^. After delivery of either scrambled control (Ctrl) or *Zbtb40* siRNA (**Fig 4d-f**) the cells were treated with osteogenic media and studied for markers of osteoblast differentiation. As these cells matured, the *Zbtb40* siRNA-treated cells exhibited a complete absence of mineralized nodule formation, as measured by Alizarin Red staining (**Fig 4g-h**). In the *Zbtb40* siRNA group, there was a reduction in staining intensity and activity (**Fig 4h-i**) of alkaline phosphatase (ALP), a marker of osteoblast differentiation. siRNA knockdown of *Zbtb40* led to reduced mRNA expression of the early transcription factor *Msx2* (**Fig 4j**), a factor that stimulates mesenchymal progenitor cell fate commitment towards the osteoblast lineage ^46^. Furthermore, there were reductions in transcript expression of the early osteoblast marker, *Col1a1*, as well as osteogenic transcription factors *Runx2* and *Sp7* (**Fig 4k**). Upon maturation, there was also a reduction in expression of *Bglap*, the gene that encodes for the mature osteoblast marker osteocalcin (**Fig 4l**).

#### Genetic manipulation of Wnt4 and Zbtb40 in vivo lead to reductions in bone mass in mice

We generated a loss-of-function mouse model to further test the effects of *Wnt4* deletion in the intact bone environment. *Wnt4* global knockout mice have been documented to die within 24 hours of birth due to disrupted kidney function ^47^. Thus, a conditional knockout mouse model was created using the Cre-Loxp system ^48^, with the Cre recombination driven by the *Prrx1* promoter to achieve targeted *Wnt4* deletion in early osteochondro-progenitors ^49^. Most bone parameters were unaffected in the *Wnt4*^*fl/fl*^ *Prrx1*-Cre mice at maturity; however, there were reductions in trabecular number at the femoral metaphysis in both female and male *Wnt4*^*fl/fl*^ femurs (**Supplemental Fig 2**). Furthermore, female *Wnt4*^*fl/fl*^ mice presented with reductions in femoral aBMD and cortical area compared to *Wnt4*^*wt/wt*^ mice (**Supplemental Fig 2**).

To further investigate the function of *Zbtb40*, a mouse model was created using the CRISPR/Cas9 technology, generating a strain (*Zbtb40*^*mut/mut*^) homozygous for a mutant or truncated form of the ZBTB40 protein. Specifically, this allele resulted in a protein product which lacked the “BTB” protein-protein interaction domain, emulating a partial loss-of-function (**Fig 5a-b**). Calvarial osteoblasts from *Zbtb40*^*mut/mut*^ mice differentiated *in vitro* show reduced ALP staining and activity as well as reduced Alizarin Red staining compared to osteoblasts from *Zbtb40*^*wt/wt*^ mice (**Fig 5c-e**). As was observed in our siRNA knockdown studies, the *Zbtb40*^*mut/mut*^ osteoblasts exhibited reduced transcript expression of *Col1a1, Runx2*, and *Sp7* early in differentiation (**Fig 5f**), as well reduced *Bglap* upon reaching maturity (**Fig 5g**). In analyzing the site-specific skeletal phenotypes of *Zbtb40*^*mut/mut*^ compared to *Zbtb40*^*wt/wt*^ mice, we found no changes in femoral areal BMD (**Fig 5h**) but did identify a significant reduction in lumbar spine areal BMD in *Zbtb40*^*mut/mut*^ male mice (**Fig 5i**). As measured by micro CT, trabecular bone in the L5 vertebrae from male *Zbtb40*^*mut/mut*^ mice exhibited reductions in bone volume/total volume (BV/TV), trabecular number (Tb.N), and connectivity density (Conn. Dens) compared to *Zbtb40*^*wt/wt*^ mice (**Fig 5j-m**). Investigating the femur in more detail, we found no differences between genotypes of either sex in cortical bone morphology at the femoral mid-diaphysis, or any differences in cortical area or thickness (**Supplemental Fig 1a-c**). Femur cortical bone strength was measured by performing three-point biomechanical bending to failure. There were no differences in maximum load or calculated stiffness (**Supplemental Fig 1d-e**) suggesting that there was no impact of this mutant allele on bone quality.

**Figure 5:**
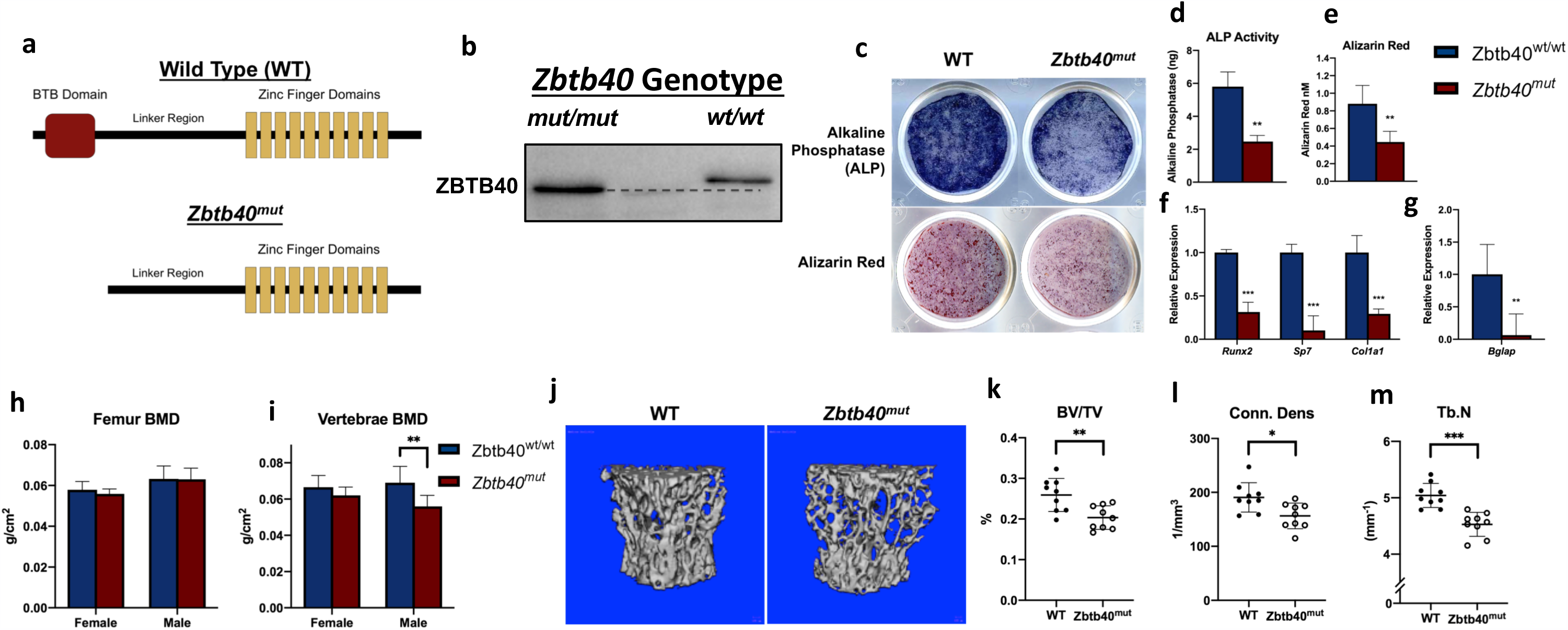
Mutation of ZBTB40 protein leads to disrupted osteogenesis and reduced vertebral bone mass in male mice: (A) Diagram of ZBTB40 protein from WT and *Zbtb40*^*mut*^ genotypes, illustrating the truncation of the BTB functional domain. (B) Immunoblot for ZBTB40 indicating the lower molecular weight of the mutant protein (∼8kDa loss). (C) Calvarial osteoblasts isolated from WT and ZBTB40^mut^ neonatal mice stained for ALP or with Alizarin Red. (D) Quantified ALP and (E) Alizarin Red values. (F) Relative mRNA levels of early and (G) late osteoblast markers taken at day 2 and day 14, respectively. (H) Bone Mineral Density (BMD) of femurs or (I) L4-L6 vertebrae from WT or ZBTB40^mut^ mice at 16 weeks of age (male wt/wt n=14, fl/fl n=9, female wt/wt n=9, fl/fl n=9). (J) 3D reconstructions and calculated (K) Bone Volume/Total Volume (BV/TV), (L) Connectivity Density (Conn. Dens), and (M) Trabecular Number (Tb.N) of trabecular bone in the vertebral body of the L5 vertebrae as measured by MicroCT (wt/wt n=9, fl/fl n=9) *\* =p<0.05 * * =p<0.01 * * * =p<0.001 -*Unpaired T test

## DISCUSSION

The traditional phenotypes used determine the genetic etiology of osteoporosis and low bone mass are BMD, which is based on X-ray attenuation by the bone tissue, or a surrogate estimate of BMD obtained via ultrasound (eBMD). Both techniques yield information about the result of a biological process, but cannot inform on what parts of that process are impacted. Thus, GWAS conducted on these phenotypes can yield loci associated with many different aspects of physiology, making experimental validation of candidate genes challenging. In this study, we chose a different route to identify loci associated with bone physiology, namely the phenotype of mineralization by the osteoblast. In this study, we established that *in vitro* osteoblast mineralized matrix formation is a heritable phenotype. To attempt to find the genetic determinants affecting osteoblast function, we identified three high resolution loci associated with the *in vitro* phenotype of osteoblast mineralization, on mouse Chrs. 1, 4 and 17. All three human regions syntenic with the mouse associations harbored associations with eBMD, suggesting that similar genes and processes (osteoblast-mediated bone formation) might be responsible.

The Chr. 1 locus, which is located at 136.6 Mb (mm9), has been previously identified in a large number of genetic mapping studies in mice ^8^. Further follow up studies using congenic mice have suggested that there are actually multiple loci on the distal end of this chromosome (ref?). Our locus overlaps with one of these loci, which is referred to in the mouse genome informatic database (MGI) as the quantitative trait locus (QTL), *bmd5* ^*50*^. This locus was found using a series of nested congenic mice for which C3H/H3J (C3H) alleles were introgressed on a C57BL/6J background. Specifically, the *c3h* alleles conferred higher volumetric BMD of the femur but had no impact on femur length ^50^. Based on the complex phenotypes of the various congenic lines, it is highly likely that this region of mouse Chr 1 contains multiple small effect size loci for BMD. No candidate gene for BMD was found for *bmd5* using the congenic approach. More recently *PTPRVP* was identified via GWAS for the phenotype of eBMD. This gene is located in the region directly homologous with our region of interest (**Fig. 3a**). Interesting, this gene is expressed at low levels in the maturing osteoblast (Gene expression omnibus series: GSE54461, ^36^), but no expression was noted in the osteoclast (Gene expression omnibus accession number: GSM1873361). Further, the mouse *Ptprv* gene is regulated by a local eQTL, suggesting that this gene could be causative for this locus.

The Chr. 17 locus interval encompasses a number of genes. Of specific interest is *Pkdcc*, as numerous GWAS in humans have identified this gene as a candidate for loci associated with whole body, femoral neck and heel ultrasound BMD ^32^. Mice lacking this gene, which is also known as *Adtk1*, present with a large number of skeletal defects including craniofacial deformities, hypo-mineralization of long bones, and limb shortening ^51^. Further, this gene is moderately expressed in the osteoblast (Gene expression omnibus series: GSE54461, ^36^). Collectively, these data imply that *Pkdcc* is a likely candidate for this Chr 17 locus.

As the candidate gene underlying the Chr 4 association remained unclear, an experimental approach was used to study this locus. The absence of an effect of *Wnt4* knockdown on *in vitro* mineralized matrix formation was an unexpected result. The WNT signaling pathway is well described in the literature to have a major role in osteogenesis and numerous WNT family members (*Wnt16, Wnt3a, Wnt5a*) ^52-54^ and canonical signaling proteins (β-catenin, LRP5/6, Dkk1/2, SOST) ^55-57^ have been shown specifically to be involved in this process. Moreover, others have shown that overexpression of *Wnt4* in osteoblasts protects against several mechanisms of bone loss ^58^; however, loss-of-function studies have been lacking. Although we did not see any effect of *Wnt4* knockdown on *in vitro* osteoblast cultures, we recognize this that cell culture system removes the osteoblast from the niche in which it normally resides. When observing the effects of Wnt4 deletion in osteoblasts *in vivo*, there were noticeable reductions in trabecular bone mass in the femur, with substantial reductions in cortical bone and aBMD in femurs of the female mice. This data conflicts with our *in vitro* data, as we do not see any changes in function of isolated osteoblasts, but observed an effect on whole bone tissue. The reason for these paradoxical changes *in vivo* may be due to the previously characterized non-canonical, paracrine effects of osteoblast-secreted *Wnt4* in altering osteoclast bone resorption through inhibiting NF-κB activation ^58^. Thus, loss of *Wnt4* in osteoblasts may not influence bone formation, but may be increasing bone resorption through cellular cross-talk, leading to the observed losses in bone mass in the femur. With the knowledge of these results, we speculate that there is likely a role for *Wnt4* as a determinant of bone mass via secretion by the osteoblast, but with the site of action being the osteoclast. While loss of *Wnt4* had no effect on intrinsic osteoblast function in isolation, loss of *Zbtb40* appears to substantially disrupt innate osteoblast gene expression, differentiation, and mineralized matrix formation. While we cannot exclude either gene (or other flanking genes for that matter), our data do suggest that *Zbtb40* is a strong candidate to mediate the effects of the osteoblast activity locus.

The *Zbtb40* gene is mostly uncharacterized, with little known about its function in any tissue. *Zbtb40* belongs to the BTB-ZF protein family, which consist of transcription factors regulating cell commitment, differentiation, and stem cell self-renewal ^59,60^. A recent study revealed that human *ZBTB40* regulates osteoblast gene transcription in cell lines, as ZBTB40 knockdown reduced mRNA expression levels of *COL1A1, RUNX2, SP7, ALP* and even *WNT4* ^61^, which is confirmatory of our results in mouse cell lines. Further, putative loss-of-function of ZBTB40 in primary cells from our mutant mouse model led to abundant reductions of mRNA expression of early osteoblast factors (*Runx2, Sp7, Msx2*). These factors have been well documented to induce commitment to the osteoblast lineage and drive osteogenic differentiation ^62-64^. Our data shows that ZBTB40 loss alters expression of these critical drivers, suggesting that, like other members of the BTB-ZF family, this gene may have an effect on cell differentiation. *Msx2*, for example, regulates early “mesenchymal” stem cell fate by stimulating osteogenesis and inhibiting adipogenesis ^46^. The fact that *Zbtb40* silencing reduces *Msx2* expression suggests that *Zbtb40* is acting upstream, which would imply a role for this gene not only in osteoblast differentiation, but at the early stage of stem cell fate determination as well. However, more investigation needs to be done to establish the function and signaling pathway of *Zbtb40*.

In summary, we have conducted a GWAS for osteoblast function *in vitro* using cells isolated from inbred strains of mice. We have shown that these loci have high concordance with human GWAS loci for eBMD. Because our phenotype was restricted to a single aspect of bone turnover, our loci become informative for the human BMD loci as we are able to provide information regarding the likely physiologic process impacted. Our data suggest that genetic studies focused on osteoblast activity have the ability to provide significant insight into the genetic and molecular basis of bone formation and osteoporosis.

## METHODS

### Animal models

All animal procedures were performed according to protocols reviewed and approved by the appropriate Institutional Animal Care and Use Committees (IACUC). All mice used in this study, with the exception of the *Zbtb40* loss of function allele mice (described in detail below), were purchased from The Jackson Laboratory. The mice were fed Laboratory Autoclavable Rodent Diet 5010 (LabDiet, Cat no. 0001326) and had *ad lib*. access to food and water. The mice were maintained on a 12hr:12hr light:dark cycle. The stock numbers for the strains used were as follows: B6;129S*-Wnt4*^*tm1.1Bhr*^*/*BhrEiJ (*Wnt4* floxed mice, #007032, 65), B6.Cg-Tg(*Prrx1*-cre)1Cjt/J (*Prrx1*-cre driver strain, # 005584), BALB/cJ (#000651), DBA/2J (#000671), FVB/NJ (#001800), RIIIS/J (#000683), BTBR_T^+^_*Itpr3*^*t*f^/J (#002282), SJL/J (#000686), NZW/LacJ (#001058), C57BL/6J (#00064), BPN/3J (#003004), TALLYHO/JngJ (#005314), NZO/HlLtJ (#002105), NON/ShiLtJ (#002423), NOR/LtJ (#002050), A/J (#000646), LG/Jm(#000675), NOD/ShiLtJ (#001976), 129S1/SvImJ (#002448), CBA/J (#000656), DDY/JclSidSeyFrkJ (#002243), AKR/J (#000648) and C3H/HeJ (#000659). The ages and genders of the animals used are described per experiment.

### Measuring mineralization in inbred strains

Primary calvarial osteoblasts were isolated from 3-9 day old neonates (males and females pooled; N=10-15 per pool) from a panel of 22 inbred strains using sequential Collagenase P digestions. Cells were plated into 6 well plates at 300,000 cells in 2ml sterile plating media (DMEM, 10% heat-inactivated FBS, 100 U/ml penicillin, 100 μg/ml streptomycin) per well. After 24 hours, confluent cells were washed 1x with DPBS (GIBCO) and placed in sterile differentiation media (MEM alpha, 10% heat inactivated FBS, 100 U/ml penicillin, 100 μg/ml streptomycin, 50 μg/ml ascorbic acid, 4 mM B-glycerophosphate). Every 48 hours thereafter cells were washed one time with DPBS (GIBCO) and differentiation media was replaced until cells were collected for analysis at day 10. Mineralized nodule formation was measured by staining cultures at 10 days post-differentiation with Alizarin Red (40 mM) (pH 5.6). The stained cells were imaged and nodule number was measured using ImageJ (NIH) ^66^. Alizarin Red was quantified by destaining cultures with 5% Perchloric acid and determining the optical density (405 nM) of the resulting solution against a standard curve. All results were obtained from three independent experiments.

### Association Analysis

To identify loci influencing mineralization, we used the Efficient Mixed Model Association (EMMA) algorithm ^35^. For the analysis, alizarin red values were rankZ transformed. SNPs were obtained from strains genotyped on the Mouse Diversity Array (http://churchill-lab.jax.org/website/MDA) ^67^. SNPs with a minor allele frequency < 0.05 were removed, leaving 196,330 SNPs. These SNPs were used to generate a kinship using the ‘emma.kinship’ R script available in the EMMA R package (available at http://mouse.cs.ucla.edu/emma/) ^35^. The emma.REML.t function of EMMA was used to perform all mapping analyses. The significance of the maximum association peak was assessed by performing 1,000 permutations of the data. In each permutation, the minimum p-value was recorded to produce an empirical distribution of minimum permutation p-values. The quantiles of this distribution were used to assign adjusted p-values. P-values exceeding a genome-wide significant of P<0.05 were used as thresholds to identify associated loci. GWAS resulted were visualized using the “qqman” R package ^68^.

### Effect of Wnt4 knockdown on mineralization

The effect upon mineralization of the *in vitro* knockdown of the *Wnt4* gene in differentiating primary calvarial osteoblasts was accomplished as outlined in Calabrese *et al* ^69^ with the following modifications. Primary calvarial osteoblast cells were isolated from the F2 generation of floxed *Wnt4* mice, plated and differentiated exactly as outlined. DNA from the tails of the 3-9 day old mice, from which the calvaria were removed, were used for genotyping via PCR using a forward primer located 5’ of the loxP site in intron 1 (GCA GAG AGG CCC AGC CTG CCC CTC A) along with a reverse primer located 3’ of the loxP site in Intron 2 (CAT GTG CCT GGC CCT AGA AAT ATC AT; expected PCR product size: 642bp (wild type allele), 819bp (floxed allele), 239bp(recombined allele)). *Wnt4* knockdown was accomplished by transfecting cells with a Cre Recombinase expression plasmid (or empty vector control, ^70^) 24h post-plating followed by the introduction of differentiation media 48h latter. Cells were collected for RNA and mineralization analysis 10 days after the initiation of differentiation as outlined except primers used for *Wnt4* expression analysis were (GAA CTG TTC CAC ACT GGA CTC (Exon2); GTC ACA GCC ACA CTT CTC CAG (Exon3). DNA was also collected at day 10 and used to qualitatively analyze the extent of Cre-recombination activity by using the genotyping primers listed above as well as quantitative analysis via qPCR with the exon2 and 3 primers. Mineral formation was measured exactly as described and the effect of *Wnt4* knockdown on mineralization was expressed as the ratio of moles of Alizarin Red bound by the Cre-transfected cells of wt/wt, wt/fl, and fl/fl genotypes.

### Cell Culture

Primary calvarial osteoblasts were cultured in the same manner as described above. The MC3T3-E1 mouse preosteoblast cell line (ATCC Subclone 4 CRL-2593) were cultured in ascorbic acid-free Alpha Minimal Essential Medium (α-MEM) supplemented with 10% Fetal Bovine Serum (FBS) and 100 U/ml penicillin/streptomycin. Cells were dissociated from the culture plate using 0.05% trypsin-EDTA and re-plated in experimental wells at 10,000 cells/cm^2^ for downstream siRNA transfection. Cells were further grown for 72 hours before reaching confluence, upon which media was changed to osteogenic medium consisting of basal culture media supplemented with 4mM β-Glycerophosphate and 50mg/ml Ascorbic Acid and differentiated until the indicated time point.

### siRNA Treatment

MC3T3-E1 cells were plated at 10,000 cells/cm^2^ in 12-well plates and allowed to adhere overnight. The following day the cells were transfected with 5nM scrambled control or *Zbtb40* Silencer Select siRNA (ThermoFisher) delivered with Lipofectamine RNAiMAX (ThermoFisher). Knockdown efficiency was determined by measuring RNA (24 hours) or protein (72 hours) expression levels post-transfection. Remaining cells were then induced with osteogenic medium at 100% confluency (72 hours post-transfection) and differentiated until the indicated time point for phenotype analysis.

### RNA Isolation and qRT-PCR Gene Expression Analysis

Cells were washed once with PBS and total mRNA was isolated using TRIzol (ThermoFisher) and purified using the GeneJET RNA Purification Kit (ThermoFisher). RNA was treated with DNAse I Amplification Grade (ThermoFisher) and reverse-transcribed into cDNA using the iScript cDNA Synthesis Kit (Bio-Rad). Real Time qPCR was performed using PerfeCTa SYBR Green FastMix (QuantaBio) and the Rotor-Gene Q Real-Time PCR Cycler (Qiagen). All reactions were performed in triplicate and normalized to *β-2-Microglobulin* using the primers F: 5’-TGACCGGCCTGTATGCTATC-3’, R: 5’-AGGCGGGTGGAACTGTGTTA-3’. Relative expression levels were calculated using the 2^-ΔΔ*CT*^ method (^71^). Primer sets: *Zbtb40* F: 5’-AGAGCCACAGCATGGAACTC-3’, R: 5’-CCGACGGAAATGGTGCAATC-3’. *Msx2* F: 5’-GATACAGGAGCCCGGCAGAT-3’, R: 5’-CTTGCGCTCCAAGGCTAGAA-3’. *Runx2* F: 5’-TGATGACACTGCCACCTCTGACTT-3’, R: 5’-ATGAAATGCTTGGGAACTGCCTGG-3’. *Sp7* F: 5’-ATGGCGTCCTCTCTGCTTGA-3’, R: 5’-CTTTGTGCCTCCTTTCCCCA-3’. *Col1a1* F: 5’-CGACCTCAAGATGTGCCACT-3’, R: 5’-GCAGTAGACCTTGATGGCGT-3’. *Bglap* F: 5’-TTCTGCTCACTCTGCTGACC-3’, R: 5’-TATTGCCCTCCTGCTTGGAC-3’.

### Western Blot

Cells were washed once with PBS then lysed and collected in Pierce RIPA buffer (ThermoFisher) supplemented with Protease/Phosphatase Inhibitor Cocktail (Cell Signaling). Lysates were vortexed three times and cleared by centrifugation (15 minutes at 15,000 rpm). Total protein was measured using the Pierce BSA Total Protein Kit (ThermoFisher) and samples were analyzed by SDS-PAGE in a 4-12% acrylamide gel (ThermoFisher). The gel was transferred to a nitrocellulose membrane using the iBlot 2 Dry Blotting System (ThermoFisher) and blocked in 5% milk in TBST for 1 hour at room temperature. The membrane was incubated overnight at 4° with Anti-ZBTB40 primary antibody (Abcam #Ab190185) or Anti-β-Actin Antibody (Sigma #A2228) and then incubated with either Goat Anti-Rabbit or Goat Anti-Mouse IgG-HRP Conjugate (Bio-rad) Secondary Antibody in 5% milk for 1 hour at room temperature. Membranes were developed using SuperSignal West Pico Sensitivity substrate (ThermoFisher) and imaged using a Bio-Rad Universal Hood II Chemiluminescent Imager. Bands were quantified using ImageJ and ZBTB40 protein levels were normalized to β-Actin.

### Cell Staining and Phenotype Measurements

At the indicated time point, cells were washed with PBS then fixed with 10% Neutral Buffered Formalin (NBF) for 10 minutes. For Alkaline Phosphatase (ALP) analysis, the fixed cells were stained with 1-Step NBT/BCIP Substrate Solution (ThermoFisher) in the dark for 30 minutes. Cells were then washed in dH_2_0 and let dry. For quantitative analysis of ALP activity, separate cells were harvested and analyzed using the SensoLyte pNPP Alkaline Phosphatase Assay Kit (AnaSpec) according to manufacturer’s instructions. To detect mineralization, cells were fixed in 10% NBF and stained with Alizarin Red for 30 minutes then washed with dH_2_0 and let dry. To obtain quantitative values for staining, plates were scanned and then de-stained with 5% Perchloric Acid. The supernatants were then collected and absorbance was measured at 405nm in a Synergy Mx Monochromator-Based Microplate Reader (Biotek).

### Zbtb40^mut^ Strain Generation

*Zbtb40*^*mut*^ mice were created on a C57BL/6J background using CRISPR/Cas9 technology, as previously described ^72^. Briefly, an insertion/deletion mutation was created on the DNA in the second coding exon of *Zbtb40* to induce a premature stop codon, limiting full translation of the generated *Zbtb40*^*mut*^ transcript. The translated ZBTB40^mut^ protein is truncated at the N-terminus by 76 amino acids, lacking a majority of the BTB domain, with the remainder of the protein intact. This has been verified by both DNA and RNA sequencing as well as western blot analysis of protein products (**Fig 5b**). This strain has been continually backcrossed to the parental C57BL/6J strain.

### Mouse Phenotyping

Dual X-ray absorptiometry (DXA) was performed on all groups using the Lunar Piximus II (GE Heathcare) as described previously ^73^. Areal BMD (aBMD), bone mineral content (BMC) and % body fat was measured at 16 weeks of age. BMD and BMC for the lumbar spine and femur was examined, as well as lean body mass. The left femur and 5^th^ lumbar vertebrae were scanned using a high-resolution micro-computed tomography system (vivaCT 40, Scanco Medical AG) to assess bone architecture. Variables computed for trabecular bone regions include: bone volume, BV/TV, and trabecular number and thickness. For cortical bone, total cross-sectional area, cortical bone area, cortical thickness, and area moments of inertia about the principal axes were computed at the femoral midshaft. Femurs were tested in 3-point flexure to failure at 0.1mm/second, collecting force and displacement data at 10Hz (Bose EnduraTec 3200, Eden Prairie, MC). Specimen alignment and orientation were co-registered accurately between imaging and mechanical test configuration. Material properties were calculated using the cortical geometric indices obtained from μCT analysis.

### Statistical analysis of Zbtb40 and Wnt4 experimental validation

Data are presented as the mean ± SD. Statistical significance of *in vitro Wnt4* experiments was determined by one-way ANOVA followed by a Dunnett’s multiple comparison’s test using *wt/wt* as the control group. Statistical significance of all other experiments were determined by Student’s unpaired *t* test. *p* values <0.05 were considered significant.

## FIGURE LEGENDS

**Supplementary Figure 1:** *Zbtb40*^*mut*^ *mice show no changes in femoral cortical bone mass or <u>strength</u>* : (A, B, C) 3D reconstructions of femur mid-diaphyseal cortical bone and calculations for cortical area and cortical thickness (wt/wt n=9, mut/mut n=9 per sex, 16 weeks of age). (D, E) Max load and stiffness calculations from biomechanical 3-point bending test (female -wt/wt n=9, mut/mut n=6, male -wt/wt n=8, mut/mut n=9, 16 weeks of age)

**Supplemental Figure 2:** *<u>Wnt4fl/fl Prrx1</u>*<u>-Cre mice show reductions in femoral trabecular number in both sexes and cortical area in females</u>: (A) Confirmation of *Wnt4* deletion by floxed allele recombination in the femur of a Cre positive *wt/fl* mouse by RT-PCR (B) Whole body and (C) Femoral BMD of female (wt/wt n=19, fl/fl n=19) and male (wt/wt n=5, fl/fl n=17) mice measured by DXA at 16 weeks of age. (D,E,F) Trabecular number (Tb.N) values and 3D reconstructions of the femoral metaphysis from female (wt/wt n=9, fl/fl n=10) and male (wt/wt n=4, fl/fl n=9) mice. (G,H) Cortical area (Ct.Ar) calculations and 3D reconstructions of the femoral mid-diaphysis in female mice. *\* =p<0.05 * * =p<0.01 * * * =p<0.001 -*Unpaired T test

