## Supplementary figures and images for "Genetic Analysis of Osteoblast Activity Identifies *Zbtb40* as a Regulator of Osteoblast Activity and Bone Mass"

### SUPPLEMENTAL FIGURES

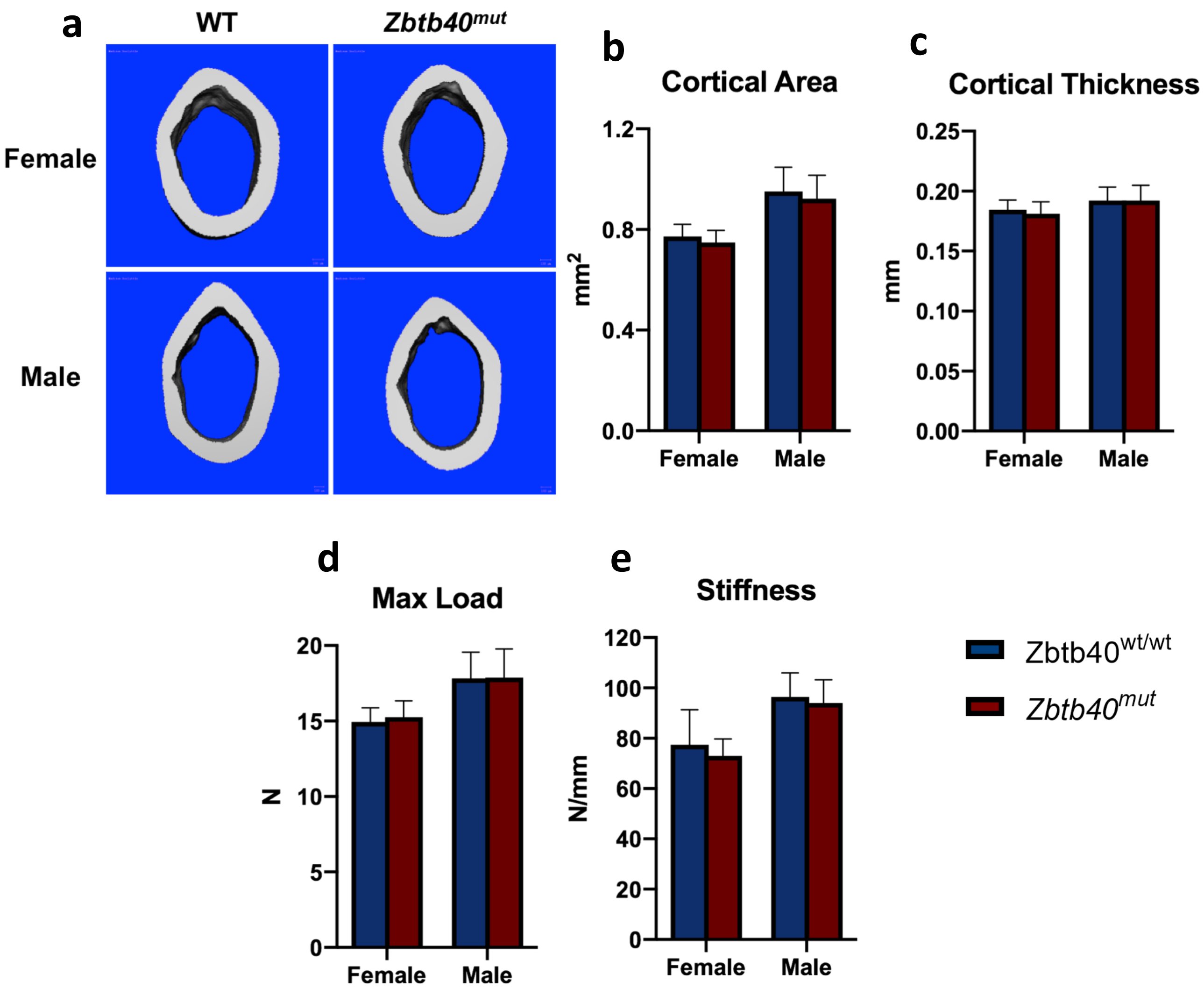

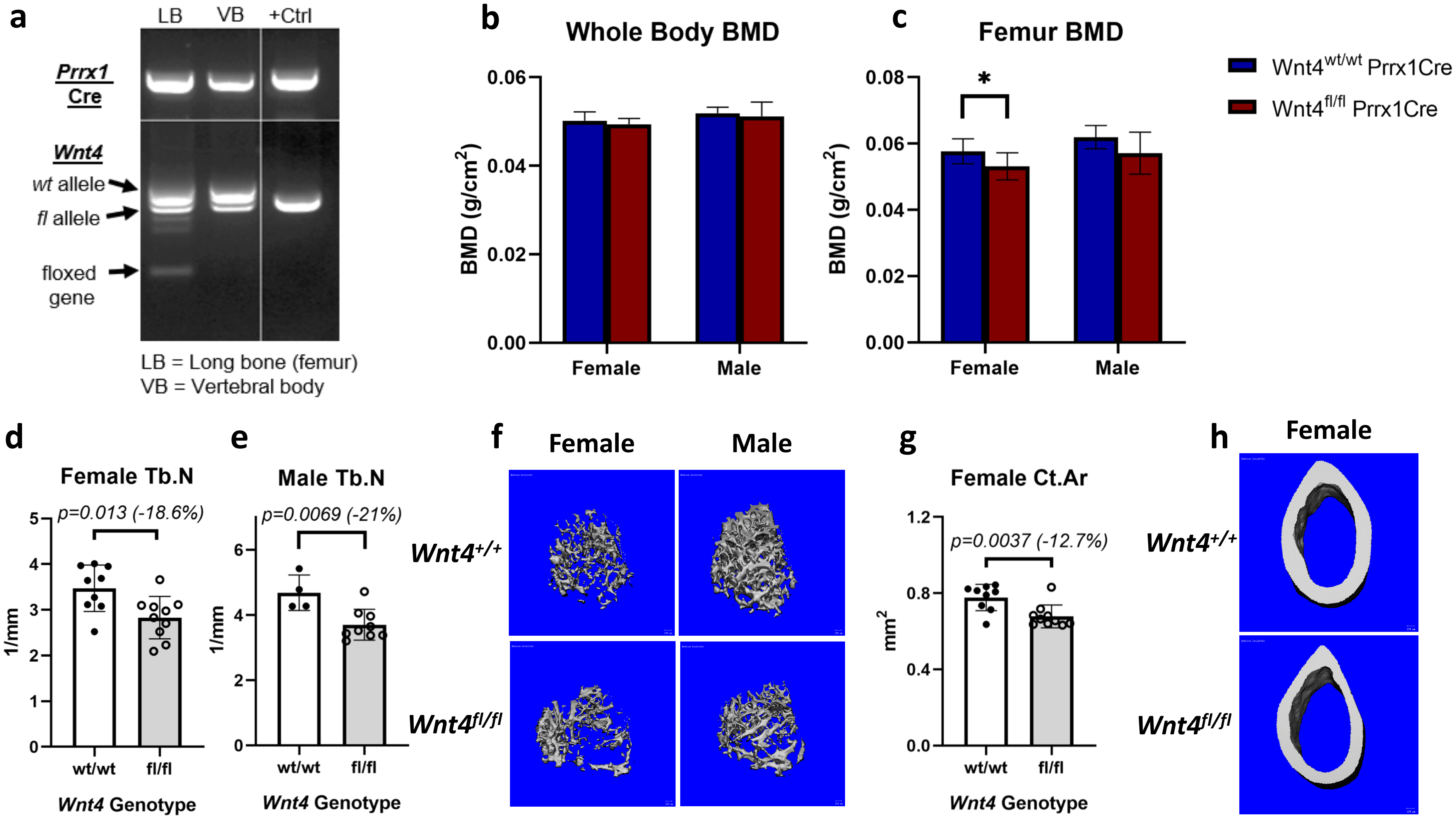
